# Network-related variations of cerebrovascular status are correlated with cognitive control performance

**DOI:** 10.64898/2026.11.19.767869

**Authors:** Xiang Quan, Kathy A. Low, Patrycja Kałamała, Frini Karayanidis, Bradley P. Sutton, Gabriele Gratton, Monica Fabiani

## Abstract

Cerebrovascular health has emerged as a crucial factor in influencing cognitive decline in aging, including cognitive control and fluid intelligence. Previous studies have largely relied on peripheral assessments of vascular health due to the paucity of global and regional estimates of cerebrovascular status. This study uses the pulse relaxation function (PReFx), a measure of pulse-wave shape based on diffuse optical tomography (pulse-DOT), to provide *regional* estimates of arterial elasticity in the cortical mantle. For the first time, we integrate a network-based approach based on previous fMRI studies and pulse-DOT regional mapping of cerebrovascular status, to test whether the support provided by the cerebrovascular system to different brain networks may differentially impact cognitive control performance. To comprehensively assess how various mechanisms of cognitive control are influenced by the cerebrovascular health of different networks, we used a task-switching paradigm that simultaneously measures proactive and reactive control processes. After removing the global signal, we found that larger positive PReFx residual values in the Dorsal Attention and Visual Networks are linked to lower cognitive control costs. Conversely, larger positive residual values in the Default Mode and Fronto-Parietal Networks are associated with higher cognitive control costs. These results indicate that the relative balance of arterial elasticity across different brain networks impacts cognitive control, highlighting the importance of studying network- based cerebrovascular status for understanding cognitive performance.

## 1. INTRODUCTION

Cerebrovascular health has emerged as a crucial factor in influencing cognitive decline in aging, including individual differences in cognitive control. In this study we integrated, for the first time, a network approach based on prior functional magnetic resonance imaging (fMRI) studies of resting state functional connectivity (rsFC; for a review see C. Gratton et al., 2018) with regional mapping of cerebral arterial status based on diffuse optical tomography (pulse-DOT; Fabiani et al., 2014) to assess how mechanisms of cognitive control may be influenced by the health of the *regional* arterial systems that perfuses different cerebral networks in younger and older adults. In the remainder of this introduction, we briefly review the main elements of this study.

### 1.1 Cognitive Control

Cognitive control involves the ability to establish and sustain relevant task goals while suppressing irrelevant distractors, and to adaptively regulate thoughts and actions to enhance task performance (G. Gratton et al., 2018). Given the central role of cognitive control among higher-order cognitive functions, extensive efforts have been devoted to developing theoretical models examining the factors that influence it (Braver et al., 2012, 2021; Miyake & Friedman 2012; Frömer et al., 2021). Based on the Dual Mechanisms of Cognitive Control (DMC) model (Braver, 2012), proactive control refers to preparatory processes that requires individuals to continuously maintain goal-relevant information before performing tasks (Miller & Cohen, 2001). In contrast, reactive control refers to immediate, responsive processes that requires individuals to adaptively address and manage stimulus-driven demands as they arise (Braver, 2012; Tang et al., 2023). The DMC framework links proactive control to sustained activation in the lateral prefrontal cortex (PFC) supported by both phasic and tonic inputs from the midbrain dopamine system, whereas reactive control involves transient activation in the lateral PFC, triggered by conflict-detection signals from anterior cingulate and medial frontal cortex (Manard et al., 2017; Braver et al., 2021; Tang et al., 2023). The differentiation between proactive and reactive control is important, as it provides strong predictions about distinct temporal dynamics and locations of brain activity that support these separable control processes.

Cognitive control processes tend to be negatively affected by aging (e.g., Braver & Barch, 2002; Xia et al., 2022) although the extent of these effects depends on many factors, including aspects of the paradigm being used (G. Gratton et al., 2018). In task-switching paradigms, aging is consistently associated with increased mixing costs, reflecting difficulty in maintaining and coordinating multiple task sets, whereas age effects on switching costs are less consistent (Karayanidis et al., 2011). In the present study we used a task-switching paradigm that differentiates cue-driven proactive control from target-driven reactive control, allowing us to examine age-related differences in both modes within the same task (Karayanidis et al., 2011; Whitson et al., 2014; Cooper et al., 2015).

### 1.2 Cerebrovascular health

The cerebrovascular system plays a crucial role in brain activity by regulating cerebral blood flow, which ensures the efficient delivery of oxygen and nutrients to the brain (Zimmerman et al., 2021). Optimal cerebrovascular functioning is essential for supporting cognitive processes, such as memory, attention and cognitive control, as it maintains the necessary blood flow to neural circuits involved in these tasks (Bowie et al., 2024; Tan et al., 2017; Jennings et al., 2008; Funke et al., 2012; Hitchcock et al., 2000; Tari et al., 2020; Leithner & Royl, 2014). Cerebrovascular dysfunction has emerged as a significant contributor to individual differences in age-related cognitive decline (Fabiani et al., 2022; Fabiani et al., 2014; Kong et al., 2020; Rabkin, 2012; Unverzagt et al., 2011; Jolly et al., 2017). This is particularly evident in vascular and other forms of dementia, where reduced cerebral perfusion and small cerebrovascular lesions can substantially impair cognitive function (van Dinther et al, 2024; Csipo et al., 2025).

Most studies investigating the relationship between cerebrovascular health and cognitive function have relied on global or peripheral measures of arterial status (e.g., carotid–femoral pulse wave velocity) rather than direct measures from the cerebral vasculature (Rabkin, 2012; Singer et al., 2014). This greatly limits our understanding of how *<u>the local cerebral vasculature</u>* impacts cognitive function. To address this gap, this study employs pulse-DOT (Fabiani et al., 2014), a diffuse optical method to examine how the cerebrovascular status of different brain network regions influences proactive and reactive cognitive control.

#### 1.2.1 Measuring cerebral arterial stiffness

Parameters of the cerebral arterial pulse waveform can be measured and mapped with pulse-DOT, which estimates arterial elasticity in the brain both globally and regionally (Fabiani et al., 2014; Chiarelli et al., 2017; Tan et al., 2017; Johnson, Ware et al., 2026; Ware et al., 2025; Mohammadi, Vincent at al., 2021). The most established and widely used measure is the Pulse Relaxation Function (PReFx; previously referred to as arterial compliance, Fabiani et al., 2014). PReFx quantifies the *shape* of the pulse waveform between the peak systole and the subsequent peak diastole, indexing how arteries return to their original size after dilating to accommodate the oxygenated blood bolus generated by a heart pulsation.

According to the Windkessel model (Westerhof et al., 2009), healthy elastic arteries have slower pulse wave velocity and slower recoil, which allows for a more distributed pulse wave during the systole and sustained blood flow during the diastole (Izzo & Shykoff, 2001). In contrast, stiffer arteries have a faster pulse wave velocity and quicker recoil, which produces a larger pulse response during the systole and a reduced ability to maintain a prolonged blood flow during the diastole. As a result, healthy arteries produce a smoother and smaller optical pulse, whereas stiffer arteries produce a larger and more “spiky” pulse. PReFx reflects the interaction between the forward pressure wave generated by the systole and the backward or recoil pressure wave from peripheral resistance (Fabiani et al., 2014). By assessing the *shape* of the arterial pulse, and particularly how well arteries retain their expanded volume after a systole, PReFx characterizes the degree of arterial elasticity in brain arteries and arterioles. The PReFx measure thus reflects a combination of arterial elasticity and peripheral resistance (Chiarelli et al., 2017).

Pulse-DOT studies have highlighted cerebrovascular status as a key factor for cognitive function and brain integrity, largely using a *global* PReFx measure (i.e., averaged across all cortical regions/voxels) (Fabiani et al., 2014; Tan et al., 2017; Bowie et al., 2024). They have consistently shown that global PReFx correlates negatively with age, and positively with gray and white matter volumes, reflecting more preserved brain structures for individuals with higher cerebrovascular elasticity (Fabiani et al., 2014; Tan et al., 2019). Global PReFx has also been found to mediate the effect of age on cognitive decline (Bowie et al., 2024; see also Tan et al., 2017; Kong et al., 2020; Clements et al., 2025). Specifically, aging has been found to negatively influence fluid but not crystallized cognition via reductions in cerebral arterial elasticity and white matter lesions (Bowie et al, 2024). Higher PReFx is also correlated with adherence to a mediterranean diet (Simpson et al., 2026). Similar optical parameters, sensitive to arterial pulsatility factors, have also shown decline in aging, with increasing cardiovascular risk factors, lower physical activity and cognition (Mohammadi, Gagnon, et al., 2021; Mohammadi et al., 2025).

A few pulse-DOT studies have also demonstrated the importance of *regional* cerebrovascular health. Chiarelli et al. (2017) showed that regional PReFx significantly explained some of the variance in brain atrophy in the corresponding regions in older adults (see also Ware et al., 2025). Fabiani et al. (2014) showed, in a double dissociation, that pulse transit time (PTT, another parameter of pulse-DOT akin to pulse wave velocity) in the left middle cerebral artery, which perfuses the left speech areas of the brain, was correlated with verbal fluency performance, while PTT in the superior portion of the precentral artery bilaterally, which perfuses the left and right prefrontal cortices, was correlated with performance in a working memory task. Tan et al. (2017) found that PReFx localized in the frontoparietal cortex predicted working memory performance more accurately than global PReFx. These findings underscore the importance of examining localized cerebrovascular health to better understand how different regions contribute to the brain’s functional organization and cognitive function.

### 1.3 Functional brain networks

The network approach, widely used in fMRI, conceptualizes the brain as a set of interconnected functional networks rather than isolated regions, allowing for a more comprehensive understanding of brain organization and cognition (Friston, 2011; Bullmore & Sporns, 2009). Applying this framework to cerebrovascular research enables the investigation of how vascular function within specific networks influences cognitive processes. This has the potential to become a powerful tool for studying cognitive-cerebrovascular interactions.

Large-scale brain networks play distinct roles in cognition and can be broadly categorized into three types: the task-positive network (TPN), the default mode network (DMN), and sensorimotor networks (Xia et al., 2022). The TPN is critical for cognitive control and attention and includes the frontoparietal (FPN), dorsal attention (DAN), and cingulo-opercular (CON) networks (Cocchi et al., 2013; Di & Biswal, 2014; Yao et al., 2020). The FPN supports conflict monitoring and resolution (Qiao et al., 2017; Yin et al., 2018), while the DAN and CON guide attentional control and salient stimulus detection (Tamber-Rosenau et al., 2018; Suo et al., 2021). In contrast, the DMN, or task-negative network, is engaged in self-reflection and internally directed thought (Anticevic et al., 2012; Raichle, 2015), typically deactivating during externally driven cognitive tasks (Harrison et al., 2008; Buckner & DiNicola, 2019). Sensorimotor networks include the visual network (VN) for early visual processing and the sensorimotor network (SMN) for motor control, and, therefore, also contribute to cognitive functions (Yeo et al., 2011; Cocchi et al., 2013; Levy & Wagner, 2011).

The organization of functional networks changes with aging, with networks becoming less distinct from each other (reduction in *segregation;* Chan et al., 2014). Kong et al. (2020) showed a strong relationship between cerebral arterial stiffness, measured globally with PReFx, and functional brain network segregation in *association networks* (i.e., CON, DAN, DMN, FPN) but not sensorimotor networks, which remained significant even when age was partialed out. Since association networks support cognitive control processes, local/regional variations in their cerebrovascular health may differentially impact cognitive performance.

### 1.4 The current study

Here we sought to examine, for the first time, the relationship between *regional* variability in arterial elasticity and cognitive control performance in healthy adults. Specifically, we examined how variability in the arterial elasticity of association networks, measured using PReFx, was associated with proactive and reactive cognitive control processes in a healthy sample spanning a broad age range (25–75 years). Based on previous findings, we hypothesized that *global* PReFx would decline with age and that higher global PReFx, signaling better arterial health, would be associated with smaller cognitive-control costs. We then examined the cortical organization of arterial elasticity across six functional networks (FPN, DMN, SMN, DAN, CON, and VN). As little is known about the network-level organization of cerebrovascular elasticity or its relationship with cognitive control performance, these analyses were considered exploratory. Specifically, we examined whether network-specific variations in PReFx residuals, after removing the global PReFx signal, formed coherent multivariate patterns and whether these variations were differentially associated with proactive and reactive control performance.

## 2. METHODS

### 2.1 Participants

Seventy-four healthy right-handed participants (age range = 25-75 years) were recruited from the Champaign-Urbana community as part of a larger, multi-session study. Eight participants were excluded from these analyses: 3 had mean accuracy or reaction time more than 3 SD from the group mean, 3 were excluded due to equipment failure during the optical recording, and 2 had average pulse measurements outside 3 SD from the mean. As such, the final sample was *N* = 66 (self-reported racial identities: 61 White, 4 Asian, and 1 Other). Age, education, sex and other characteristics of the final sample are presented in **Table 1**. All participants included in the study were native English speakers and reported themselves to be in good health with normal hearing, normal or corrected-to-normal vision, and free from medications specifically intended to affect the central nervous system.

**Table 1.**
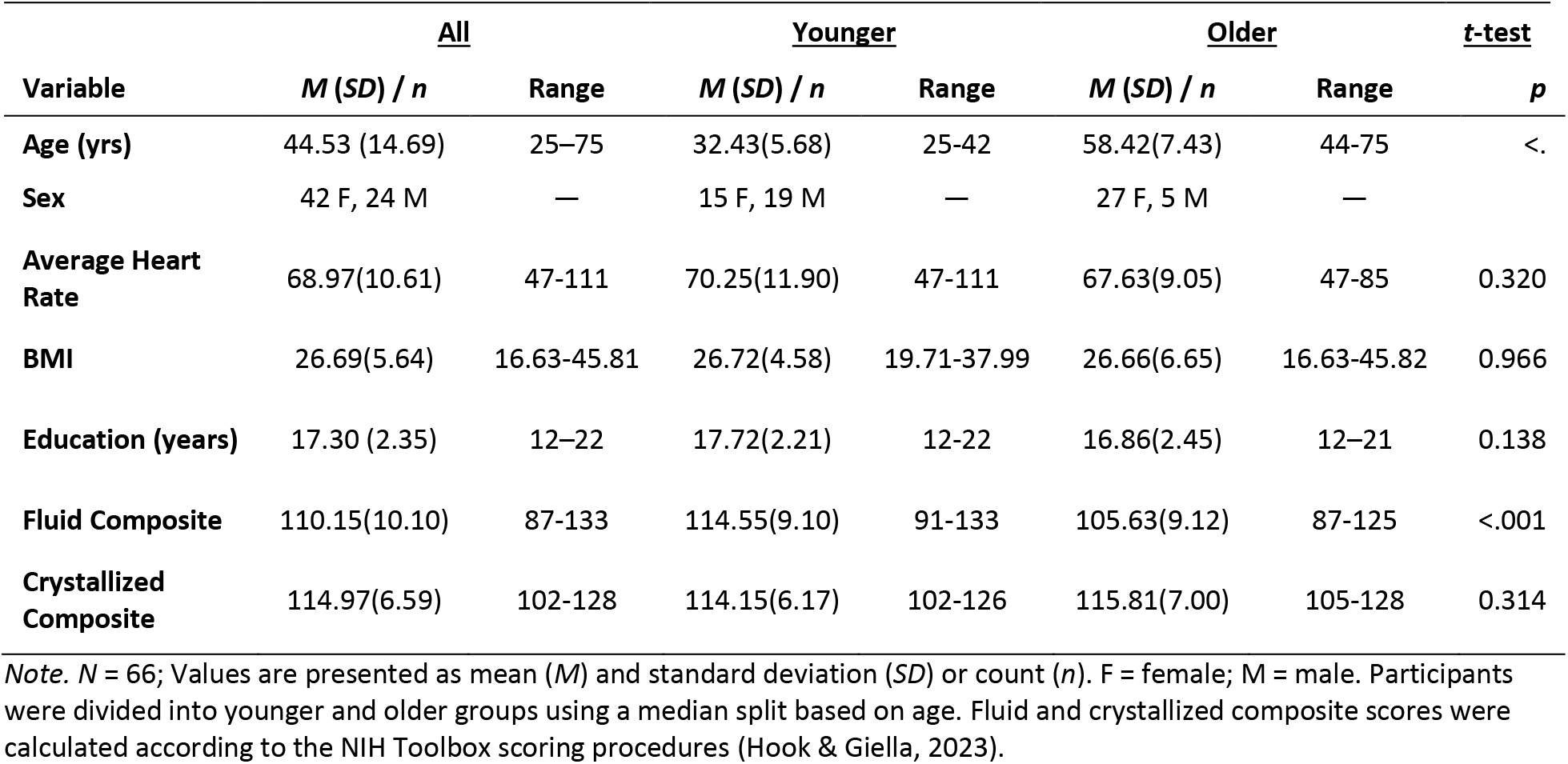
Sample Characteristics.

|  | **All** |  | **Younger** |  | **Older** |  | ***t*-test** |
|---|---|---|---|---|---|---|---|
| **Variable** | ***M* (*SD*) / *n*** | **Range** | ***M* (*SD*) / *n*** | **Range** | ***M* (*SD*) / *n*** | **Range** | ***p*** |
| **Age (yrs)** | 44.53 (14.69) | 25–75 | 32.43(5.68) | 25-42 | 58.42(7.43) | 44-75 | <. |
| **Sex** | 42 F, 24 M | — | 15 F, 19 M | — | 27 F, 5 M | — |  |
| **Average Heart Rate** | 68.97(10.61) | 47-111 | 70.25(11.90) | 47-111 | 67.63(9.05) | 47-85 | 0.320 |
| **BMI** | 26.69(5.64) | 16.63-45.81 | 26.72(4.58) | 19.71-37.99 | 26.66(6.65) | 16.63-45.82 | 0.966 |
| **Education (years)** | 17.30 (2.35) | 12–22 | 17.72(2.21) | 12-22 | 16.86(2.45) | 12–21 | 0.138 |
| **Fluid Composite** | 110.15(10.10) | 87-133 | 114.55(9.10) | 91-133 | 105.63(9.12) | 87-125 | <.001 |
| **Crystallized Composite** | 114.97(6.59) | 102-128 | 114.15(6.17) | 102-126 | 115.81(7.00) | 105-128 | 0.314 |

*Note.* *N* = 66; Values are presented as mean (*M*) and standard deviation (*SD*) or count (*n*). F = female; M = male. Participants were divided into younger and older groups using a median split based on age. Fluid and crystallized composite scores were calculated according to the NIH Toolbox scoring procedures (Hook & Giella, 2023).

Participants also reported no history of neurological or psychiatric disorders and had no signs of dementia (as assessed by the Montreal Cognitive Assessment; range of this sample: 24-30; Nasreddine et al., 2005) or depression (as assessed by the Beck’s Depression Inventory; range of this sample: 0-17; Beck et al., 1996). All participants signed informed consent, and all procedures were approved by the Institutional Review Board of the University of Illinois Urbana-Champaign.

### 2.2 Task-switching paradigm

To assess cognitive control, participants performed a task-switching paradigm (Whitson et al., 2014). The paradigm comprised two choice tasks (**Fig. 1**): a Letter classification task (i.e., vowel: A, E, I, U, or consonant: G, K, N, R) and a Number classification task (i.e., odd: 3, 5, 7, 9 or even: 2, 4, 6, 8). The Letter and Number tasks were mapped to Cold and Hot color cues, respectively. Vowel/odd and consonant/even responses were mapped to left and right index finger responses, respectively. The cue color was never repeated on successive trials to eliminate any confounding effect of cue identity repetition (Logan & Bundesen, 2003). Examples of color-task and response-task mappings are shown in **Fig. 1**.

**Figure 1.**
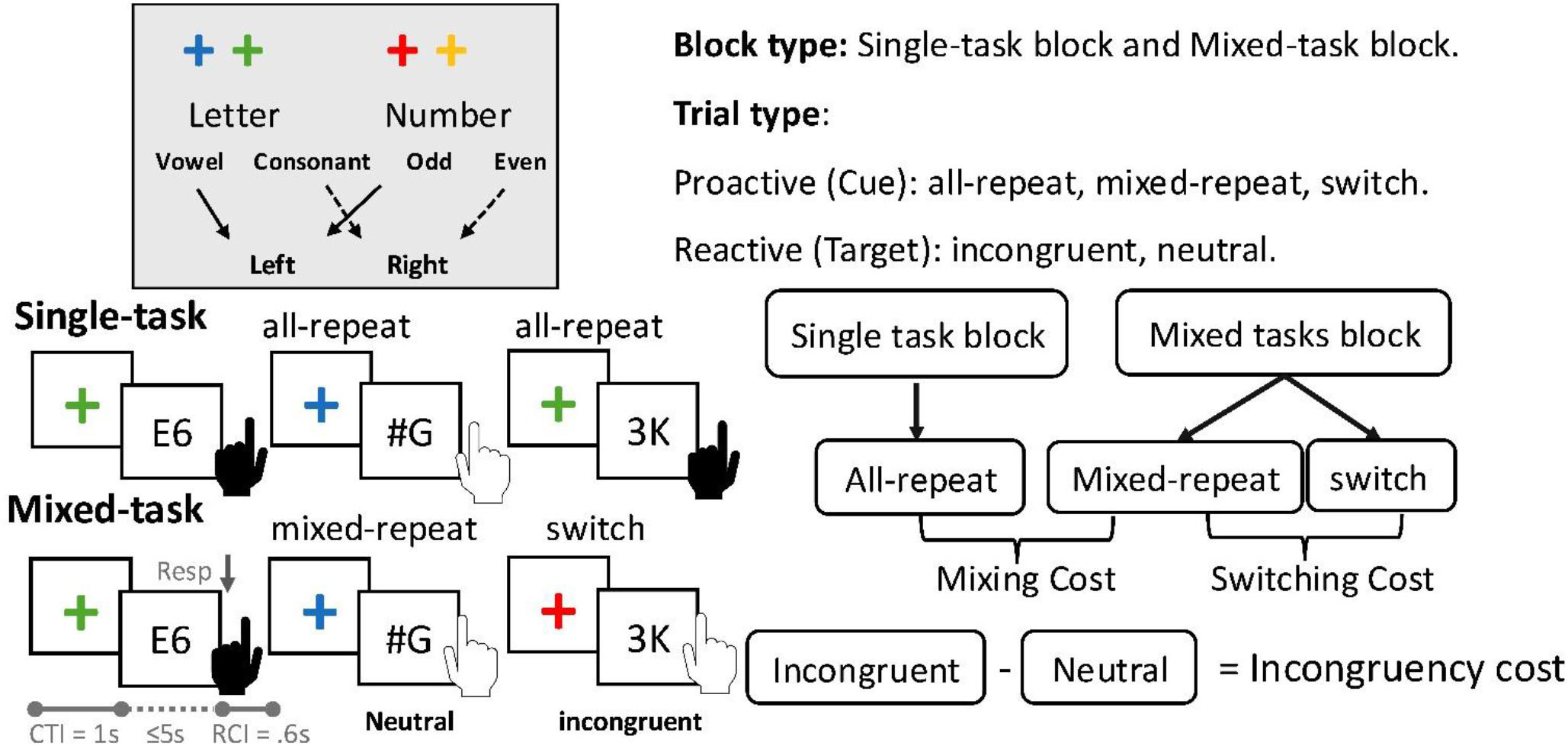
Experimental design. Upper left panel: Schematic of the task-switching paradigm. Cold colors (blue or green) cue the letter classification task while hot colors (red or orange) cue the number classification task. Participants respond based on the task relevant stimulus while ignoring the task-irrelevant stimulus. Each classification task is mapped onto opposite index fingers. Bottom left panel: Example trials: When two consecutive trials cue the same task, they are classified as repeat trials; when they cue different tasks, they are classified as switch trials. Right panel: Block and trial types. All-repeat trials occur in single-task blocks, whereas mixed-repeat and switch trials occur in mixed-task blocks. Mixing cost (mixed-repeat − all-repeat) and switching cost (switch − mixed-repeat) index proactive control, whereas incongruency cost (incongruent − neutral) indexes reactive control. CTI: cue-trial interval.

A trial consisted of a task cue (color cross) that validly indicated the relevant task and was replaced after 1,000 ms by the target stimulus (cue-target interval, CTI) which was presented until a response or 5,000 ms elapsed. The next trial began 600 ms after the response (response-cue interval, RCI = 600 ms; response-target interval, RTI = 1,600 ms). Incorrect responses were followed by an error feedback tone, and the following trial was excluded from all analyses.

The target consisted of two characters. One character was selected from the task-relevant stimulus set (e.g., a letter when the Letter task was cued; see top left panel of **Fig. 1**). On 50% of trials, the other character was selected from the task-irrelevant stimulus set (e.g., a number when the Letter task was cued) and was mapped to the opposite hand response to the target (e.g., if target was *K* that was mapped to a right-hand response, the task-irrelevant character was an odd number mapped onto a left-hand response). These trials are referred to as **incongruent targets**. On the remaining 50% of trials, the other character was selected from a set of non-alphanumeric symbols (i.e., %, #, ?) that were not mapped to any response; these are referred to as **neutral targets**. The letter-number pairs for incongruent targets and letter-symbol or number-symbol pairs for neutral targets were randomly selected on each block of trials from the full set of paired exemplars, with the restriction that the same exemplar could not be repeated on successive trials.

Mixed-task blocks included a unique pseudorandomized sequence of equal number of mixed-repeat trials (e.g., a blue cue on trial n-1 was followed by a green cue on trial n, indicating that the Letter task would be repeated) and switch trials (e.g., a green cue on trial n-1 was followed by an orange cue on trial n, indicating a switch from Letter to Number task), with no more than four successive repeat or switch trials. Single-task blocks required repeating the same task on all trials; i.e., they included only cold (Letter task) or hot (Number task) cues but included both neutral and incongruent targets. To differentiate between repeat trials in single-task and mixed-task blocks, we refer to them as all-repeat and mixed-repeat trials, respectively.

The task-switching paradigm included ten blocks of 82 trials each, comprising four single-task blocks (two Letter and two Number task blocks) and six mixed-task blocks presented in a fixed order (alternating sequence of pairs of mixed-task and single-task blocks (Number then Letter). This yielded 246 mixed-repeat, 246 switch trials and 328 all-repeat trials, each with approximately equal distribution of task (letter, number) and target type (neutral, incongruent). Participants were instructed to respond as quickly and as accurately as possible. Following each block, average reaction time and accuracy feedback was displayed and participants were encouraged to monitor and improve their performance based on this feedback.

#### 2.2.1 Practice sessions

Before completing the task-switching paradigm, participants completed two short practice sessions approximately 1-2 weeks apart. The first practice session included 10 short single-task and mixed-task blocks (20-40 trials) with varying cue-trial intervals (CTI, 1000 or 2000 ms), and response time limits (unlimited or 5,000 ms) to develop cue-task-response associations. The second practice session was administered immediately before the task-switching experiment and on the same day as the optical data collection. It consisted of four blocks: a single-task blocks for the Number and Letter tasks (20 trials each), followed by two mixed-task blocks (40 trials each), all with the same CTI (1000 ms) and response time limit (5000 ms) as the main task. If the switch trial accuracy in the mixed-task blocks was below 50%, participants completed two additional mixed-task practice blocks before proceeding.

#### 2.4 Data Acquisition

The pulse-DOT data were collected during two resting-state optical recording blocks, concurrently with the EKG data, and were followed by the collection of the behavioral task-switching data. Within the same session, the locations of the optodes were also digitized (see 2.7.1). The structural MRI data were recorded in a separate session. Data were collected as part of a larger study and data collection for any given participant was completed within two weeks.

#### 2.4.1 Resting-state optical recording

Resting-state optical data were acquired with 12 synchronized, multichannel frequency-domain oximeters (ISS Imagent®, Champaign, Illinois) with 256 laser diodes (128 emitting light at 690 nm and 128 at 830 nm) as sources and 48 photo-multiplier tubes (PMTs) as detectors. Two 10-minute resting-state blocks were recorded while participants were comfortably seated and were instructed to remain still and keep their eyes open.

Time-division multiplexing was employed so that each detector picked up light from 16 different sources at different times within a multiplexing cycle with a final sampling rate of 39.0625 Hz. The light was transmitted to the scalp using single optic fibers (0.4-mm core) and from the scalp back to the photomultiplier tubes using optic fiber bundles (3-mm diameter). The fibers were held in place using soft, semirigid, custom-built helmets, fitted to participants based on their head circumference.

The pulse-DOT measurements were based on a high-density optode montage, consisting of 48 detectors each coupled with 16 time-multiplexed sources (half at 830 nm and half at 690 nm), resulting in a total of 768 channels. As can be seen in top panel of **Fig. 2**, which shows the optode locations on a structural MRI, this montage provides broad coverage of most of the cortical mantle. The bottom panel of this highlights the area of optical sensitivity.

**Figure 2.**
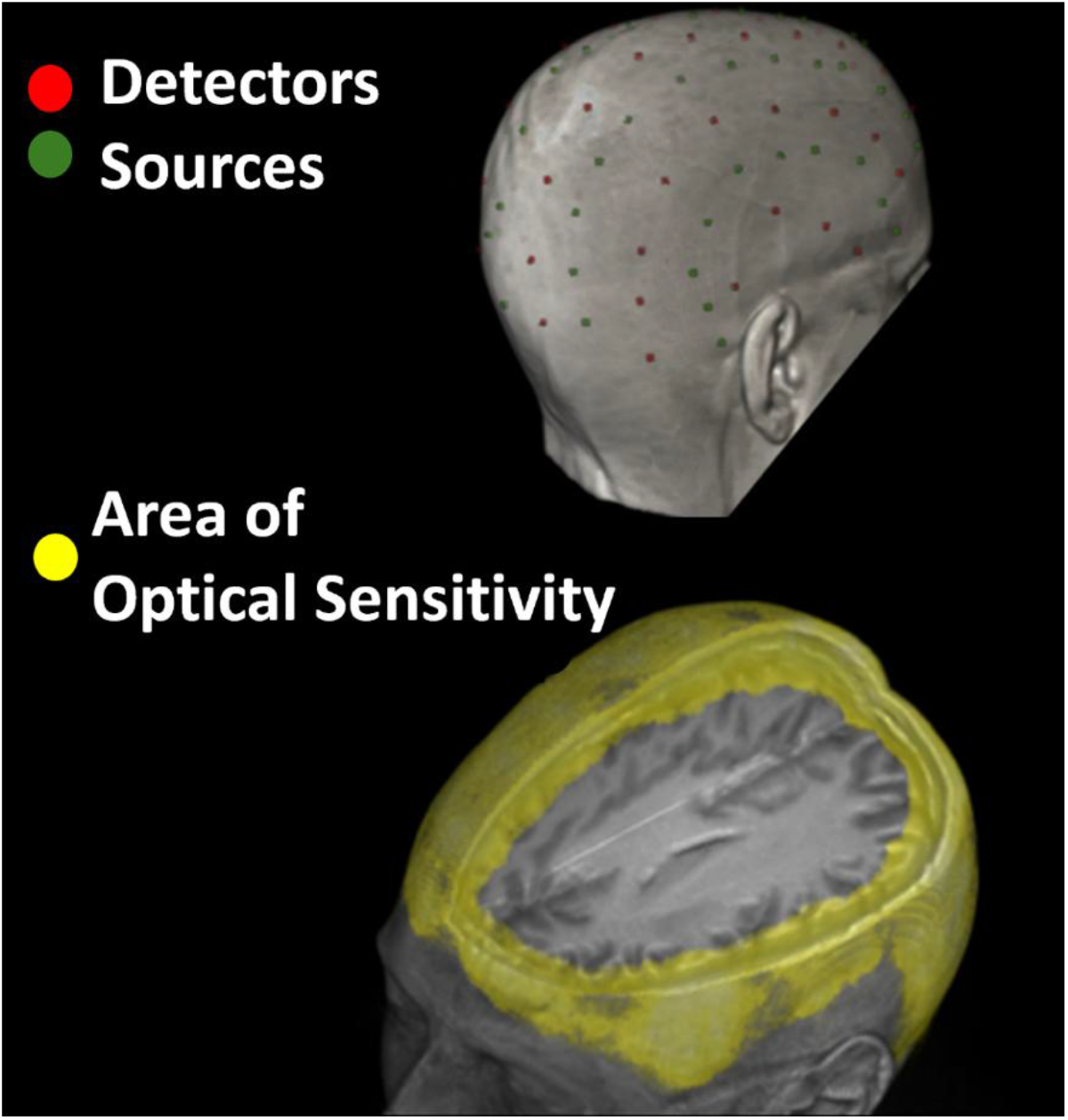
**Top**. Source and detector locations for pulse-DOT recording displayed on the MRI of a representative participant. Red dots represent the locations of the detectors and green dots those of the sources. Only the sources and detectors visible from the right posterior view are shown here, but the montage used was symmetric and included similar sources and detectors in locations not visible from this view. **Bottom.** The yellow area represents the area of optical sensitivity, also symmetrical on all views.

#### 2.4.2 EKG recording

Concurrently with pulse-DOT data acquisition, we recorded the electrocardiogram (EKG), using Brain Vision Recorder software and a Brain Vision V-Amp (Brain Products®, Germany). Concurrent and time-locked acquisition allowed us to synchronize the optical pulse data to the R wave of the EKG and ensure that the same pulse was examined regardless of recording location. Specifically, lead 1 of the EKG (left wrist referenced to right wrist) was recorded with a sampling rate of 200 Hz and a band-pass filter of 0.1– 100 Hz, with a 60 Hz notch filter. The exact timing of each R-wave peak was determined by searching for peak points exceeding a voltage threshold (scaled for each participant) and dismissing any peak points outside the normal range of inter-beat intervals. The identification of each peak was verified by visual inspection, and false detections were eliminated.

#### 2.4.3 Structural MRI

A T1-weighted, structural magnetic resonance image (sMRI) was obtained for each participant with 3T Siemens Prisma MR whole-body scanner (Siemens, Munich, Germany) using a 20-channel head coil. An MPRAGE image was acquired with the following parameters: voxel size = 0.8 x 0.8 x 0.8 mm, field-of-view = 179.2 x 256 x 256 mm, acceleration factor = 2; slice thickness = 0.80 mm; 224 sagittal slices; FA = 8°; TR = 2,400 ms; TI = 1,060 ms; TE = 2.31 ms.

#### 2.4.4 Co-registration of optode locations with structural MRI

The spatial locations of individual optical sources and detectors (see **Fig. 2**) were digitized using a Polhemus “3Space” FASTRAK 3D digitizer (Polhemus, Colchester, VT) using a recording stylus and three head-mounted receivers, which allowed for small movements of the head during measurement. The location of the nasion and preauricular points served as reference points. The Polhemus digitization points were then co-registered with the sMRI using the three fiducial markers and/or a surface-fitting and scalp-forcing procedure (Chiarelli et al., 2015; see also Whalen, et al., 2008). The individually coregistered data were then transformed into MNI space to permit registration across participants (Evans et al., 1992), as shown in **Fig. 2**.

### 2.5 Data processing and analysis

#### 2.5.1 Behavioral data

As shown in the right panel of **Fig. 1**, *mixing cost* was quantified as the difference between mixed-repeat and all-repeat trials, whereas *switching cost* was measured by the difference between switch and mixed-repeat trials (Karayanidis et al., 2011). These two measures were used as indices of <u>proactive control</u>, reflecting the additional demands associated with maintaining task goals. *Incongruency cost* was quantified as the difference between incongruent and neutral trials and was used as an index of <u>reactive control</u>, reflecting the additional processing required to resolve conflict during target interval. We conducted three separate linear mixed-effects models to examine proactive control (mixing cost and switching cost) and reactive control (incongruency cost). For proactive control, two separate linear mixed-effects models were fitted to examine mixing and switching costs, respectively. The mixing cost model included all-repeat and mixed-repeat trials, whereas the switching cost model included mixed-repeat and switch trials. For both models, we averaged across Target Type (neutral, incongruent) within each Trial Type. In each model, trial type was specified as a two-level within-subject categorical factor using difference coding (−0.5, +0.5), such that the Trial Type coefficient represented the difference between the two conditions and the Age coefficient represented the effect of age averaged across conditions. Age was mean-centered and entered as a continuous predictor, and the Trial Type × Age interaction was included. Subject was included as a random intercept. For reactive control, a separate mixed-effects model was fitted in which congruency was specified as a two-level within-subject categorical factor comprising neutral and incongruent trials and was also difference-coded (−0.5, +0.5). For this model, we averaged across Trial Type (all-repeat, mixed-repeat, and switch) within each Target Type. The model included mean-centered Age, the Target Type × Age interaction, and a random intercept for Subject. Fixed effects were evaluated using t-tests with Satterthwaite’s approximation for degrees of freedom.

Models were fitted using the *lme4* package (Bates et al., 2015), *lmerTest* (Kuznetsova et al., 2017) and *emmeans* (Lenth, 2024). All analyses were conducted in R (R Core Team, 2024) using RStudio (Posit Team, 2024). The inverse efficiency score (IES = Reaction Time/Accuracy) quantifies the time participants require to complete a task, adjusted for their accuracy (Bruyer & Brysbaert, 2011; Vandierendonck, 2018). When there is a trade-off between speed and accuracy, it allows to compensate for differences in accuracy, providing a robust measure of performance (Townsend & Ashby, 1978). Because tradeoffs between speed and accuracy often differ between younger and older adults, we used IES to avoid bias in this broad age-range sample. Here we only report the IES results as they were largely consistent with those or RT and accuracy. The RT and accuracy results are provided in the Supplementary Materials.

#### 2.5.2 Pulse-DOT data

The optical AC intensity data were normalized and low pass filtered at 10 Hz using a Butterworth filter. In-house software Opt-3D (Gratton (2000) was used to reconstruct voxel-level optical signals by combining data from channels whose diffusion paths intersected each voxel. Only channels with source-detector distances between 2 and 5.5 cm were included to retain adequate sensitivity to cortical signal while minimizing contamination from superficial tissues and noise. The voxel data were then surface projected (2D model) to the superior axial, left/right sagittal, and anterior and posterior coronal views. For each channel, each voxel was assigned a weight related to the estimated sensitivity of the measures to phenomena (absorption and scattering) occurring in brain regions underlying that voxel. This sensitivity was estimated based on a perturbation model presented by Feng and colleagues (1995). For each surface voxel, the intensity data were sorted in epochs time-locked to each R-wave from the EKG to create single trial pulse waveforms, which were then baseline-corrected by subtracting the mean signal measured 154-256 ms after the R peak, corresponding to the first post-R-peak diastole interval. PReFx was computed as the area under the pulse wave between the peak systole and the following peak diastole, normalized for inter-beat interval and peak amplitude to avoid confounds due to pulse magnitude and cardiac-cycle duration. A constant of 0.5 was then subtracted from the normalized area. This value represents a hypothetical linear relaxation trajectory, in which the arterial wall relaxes at a constant rate between peak systole and the subsequent diastolic minimum. PReFx was computed for each individual pulse cycle and voxel. PReFx values outside the valid range of 0-1 were considered outliers and discarded. Similarly, the data for a pulse cycle and voxel were discarded if the pulse amplitude was >3 dB or smaller than .1 dB, and if the inter-beat interval estimated from the optical data differed by more than 76.8 ms (three samples at 39.0625 Hz) from that estimated from the EKG (this typically occurred when movement artifacts significantly altered the optical data). To avoid estimating pulse parameters from insufficient data, voxels with fewer than 10% of the trials meeting criteria were excluded from further analyses. For each voxel, PReFx values were averaged for all acceptable trials/pulsations.

##### 2.5.2.1. PReFx calculation: Global, and network specific computations

**Fig. S1** in the Supplementary Materials presents a detailed workflow for calculating PReFx for the global and network analyses. *Global PReFx* was computed by averaging the PReFx values of all acceptable voxels. To assess *PReFx in individual networks*, we created ROI masks (**Fig. S2**) based on a cortical surface parcellation atlas developed by Yeo and colleagues (2011). The original parcellation included seven networks. However, the recording of optical signals is restricted to regions of the cortex close to the surface of the head, typically within 3 cm (Chiarelli et al., 2016; Gratton & Fabiani, 2010; see **Fig. 2**). Therefore, we excluded the limbic network ROI mask due to its deep position within the brain and only included six ROIs in the DMN, FPN, CON, SMN, DAN, and VN networks that were located less than 3 cm from the scalp. Next, we averaged the voxels values for each ROI network mask separately, to obtain the average PReFx value for each ROI network mask within each subject (raw network PReFx).

Finally, we computed the **deviations** or **residuals** of each ROI PReFx value from the global PReFx value to determine whether any given ROI had a PReFx value greater or smaller than the global PReFx (see **Fig. S1**). This method is conceptually akin to the fMRI removal of the global signal in network analysis and enhances sensitivity to network-specific variations by isolating the relative arterial health of each network in the context of overall brain arterial health (Macey et al., 2004; Desjardins et al., 2001). By subtracting the global PReFx, we effectively removed shared influences, thereby clarifying the unique contributions of individual networks. This approach also improves the interpretability of the results by highlighting how residuals in network’s arterial health from the global signal are associated with cognitive control performance.

##### 2.5.3.1 Principal Component Analysis

To understand the underlying structure and relationships among different networks, we used a Principal Component Analysis (PCA; *psych* package in R; Revelle, 2017) to investigate how PReFx residual scores from the six neural networks cluster together. The analysis included PReFx residual scores from DMN, FPN, CON, DAN, SMN, and VN. To facilitate the interpretation of the principal components, a varimax rotation was applied to maximize the independence of factors and enhance the clarity of the underlying factor structure. The number of principal components retained was determined by visual inspection of the scree plot using the elbow rule (Jolliffe, 2002) and consideration of the cumulative variance explained.

#### 2.5.4 Partial Least Squares Correlation

To explore the latent relationships between age, sex, the PReFx residual scores from the six neural networks and cognitive control performance, while controlling for multiple comparisons, a Partial Least Squares Correlation (PLSC) analysis was performed using *TExPosition* package (Beaton et al., 2014). Specifically, this analysis involved two sets of variables: independent variable (*X*) block included age, sex (male = 0, female = 1) and PReFx residual scores. Dependent variable (*Y*) block included IES mixing cost, switching cost and incongruency cost. All variables besides sex in both blocks were mean-centered and standardized to unit variance. PLSC was used to identify latent structures and correlations between *X* and *Y* blocks, revealing how variations in PReFx scores and demographic factors relate to cognitive control performance.

Statistical significance of the latent dimensions was assessed using 1000 permutations. For each permutation, the participant labels of one data block were randomly permuted while the other block was held fixed, thereby disrupting the correspondence between the X and Y blocks while preserving the multivariate structure within each block. Permutation-based *p* values for the eigenvalues were used to determine which latent dimensions explained greater cross-block covariance than expected under the null hypothesis.

Signed variable contributions were used descriptively to characterize the direction and relative contribution of each observed variable to the latent dimensions. The stability of these variable contributions was evaluated using 1000 bootstrap resamples. Participants were resampled with replacement, with the same resampling indices applied to both the X and Y blocks to preserve their correspondence. Bootstrap ratios (BSRs) were calculated as the bootstrap mean contribution divided by its bootstrap-estimated standard deviation; variables with |BSR| > 2 were considered to show stable contributions to the latent dimension.

## 3. RESULTS

### 3.1 Behavior

Three separate linear mixed-effects models were fitted to subject-level condition-averaged IES values to examine proactive control (mixing cost and switching cost) and reactive control (incongruency cost). The estimated effects of trial type, target type, age, and their interaction were evaluated using *t*-tests with Satterthwaite’s approximation for degrees of freedom (**Table 2**) (Kuznetsova et al., 2017). **Fig. 3** summarizes the mixing, switching, and incongruency effects.

**Figure 3.**
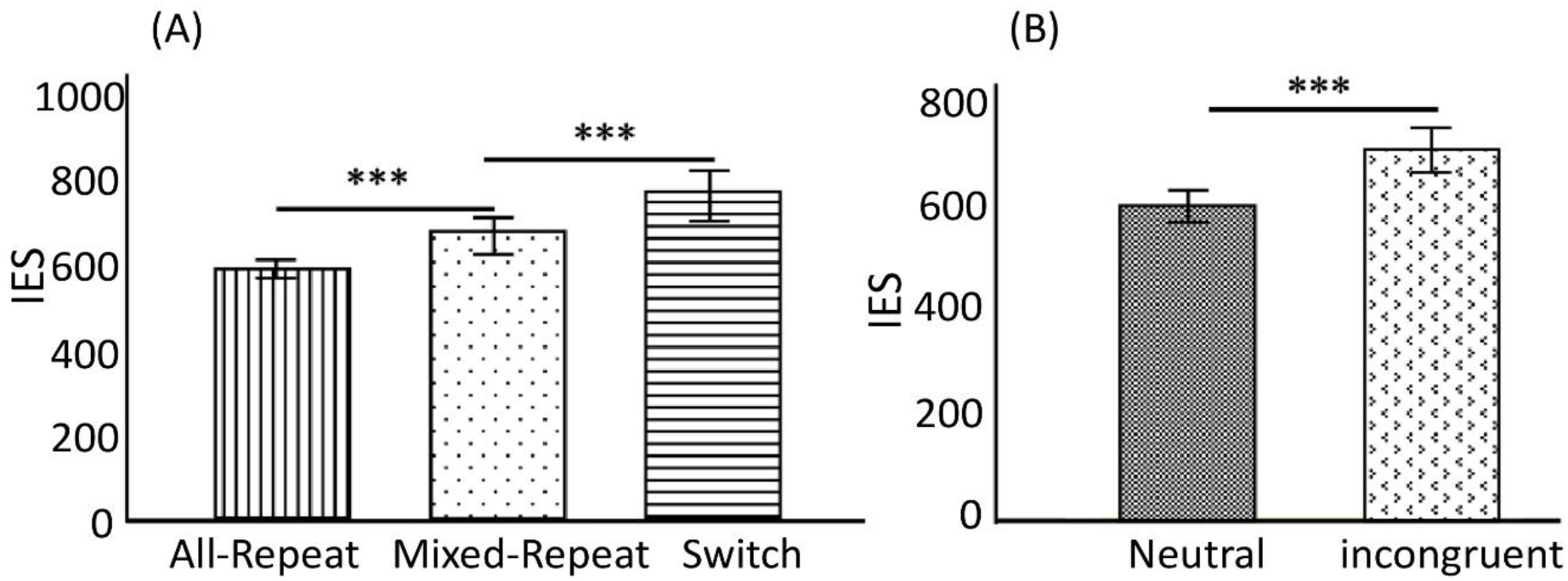
Estimated marginal means of the inverse efficiency scores (IES) across conditions. (A) IES for all-repeat, mixed-repeat, and switch trials; the all-repeat vs. mixed-repeat and mixed-repeat vs. switch differences correspond to mixing and switching costs, respectively. (B) IES for neutral and incongruent trials; their difference corresponds to the incongruency cost. Error bars represent 95% confidence intervals. Significance markers indicate the corresponding condition contrasts reported in the linear mixed-effects models. \*\*\**p* < .001.

**Table 2.** Linear Mixed-Effects Model Results for IES across Mixing, Switching, and Incongruency Costs with Age (years) as a predictor.

| **Outcome** | **Effect** | **Estimate** | **SE** | **df** | ***t*** | ***p*** |
|---|---|---|---|---|---|---|
| Mixing cost | **Dependent variable: IES (RT / ACC)** |  |  |  |  |  |
|  | Age | 2.44 | 1.04 | 64 | 2.34 | 0.022<sup>*</sup> |
|  | Trial type (all-repeat, mixed-repeat) | 86.03 | 11.92 | 64 | 7.22 | < 0.001<sup>***</sup> |
|  | Age × Trial type | 0.054 | 0.82 | 64 | 0.07 | 0.948 |
| Switching cost | **Dependent variable: IES (RT / ACC)** |  |  |  |  |  |
|  | Age | 2.23 | 1.67 | 64 | 1.33 | 0.187 |
|  | Trial type (mixed-repeat, switch) | 89.52 | 12.59 | 64 | 7.11 | < 0.001<sup>***</sup> |
|  | Age × Trial type | -0.48 | 0.86 | 64 | -0.55 | 0.583 |
| Incongr. cost | **Dependent variable: IES (RT / ACC)** |  |  |  |  |  |
|  | Age | 2.35 | 1.24 | 64 | 1.89 | 0.063. |
|  | Target type (Neutral, Incongruent) | 107.73 | 10.18 | 64 | 10.58 | < 0.001<sup>***</sup> |
|  | Age × Target type | 0.51 | 0.70 | 64 | 0.72 | 0.472 |

*Note.* IES = inverse efficiency score; RT = reaction time (ms); ACC = accuracy; SE = standard error; *df* = degrees of freedom. Incongr. = incongruency. Degrees of freedom was approximated using Satterthwaite’s method. *IES* = inverse efficiency score. <sup>\*</sup> *p* < .05; <sup>\*\*</sup> *p* <.01; <sup>\*\*\*</sup> *p*<0.001

As shown in **Table 2**, the proactive control analyses revealed significant mixing and switching costs, with higher IES for mixed-repeat than all-repeat and for switch than mixed-repeat conditions, respectively. IES increased significantly with age when averaged across conditions in both the mixing and switching cost models. These findings replicated previous behavioral results for mixing and switching costs (Karayanidis et al., 2011). However, neither the age × mixing cost nor the age × switching cost interactions were significant, providing no evidence that either cost varied with age. For reactive control, there was a significant incongruency cost, with higher IES for incongruent than neutral trials. IES also increased significantly with age when averaged across neutral and incongruent conditions, whereas the Age × Incongruency cost interaction was not significant.

Thus, although behavioral performance showed a main effect of age, there was no evidence that the magnitude of the mixing, switching, or incongruency costs interacted with age (**Table 2**). The RT and Accuracy results are provided in Supplementary Materials **Table S1**.**3.2 Correlation Results.** We calculated three types of correlations. First, we examined correlations between global PReFx, age, and behavior, to check for consistency with previous findings. Next, we calculated the correlations between the raw network ROI PReFx and behavior, to verify the need to remove the global signal as done in studies of rsFC (see **Fig. S1**). Finally, as planned, we calculated the correlations between residual network-related PReFx and indices of IES costs derived from the task switching paradigm. Results for each of these are presented below.

#### 3.2.1 Global PReFx

As shown in **Table 3**, age was significantly correlated with global PReFx, indicating that aging was associated with reduced arterial elasticity, replicating previous studies (e.g., Fabiani et al., 2014; Tan et al., 2019; Bowie et al., 2024). Global PReFx was also negatively correlated with incongruency cost, even after controlling for age and sex (*r* = -0.261, *p* = 0.037), suggesting that better arterial elasticity is associated with greater reactive control.

**Table 3.**
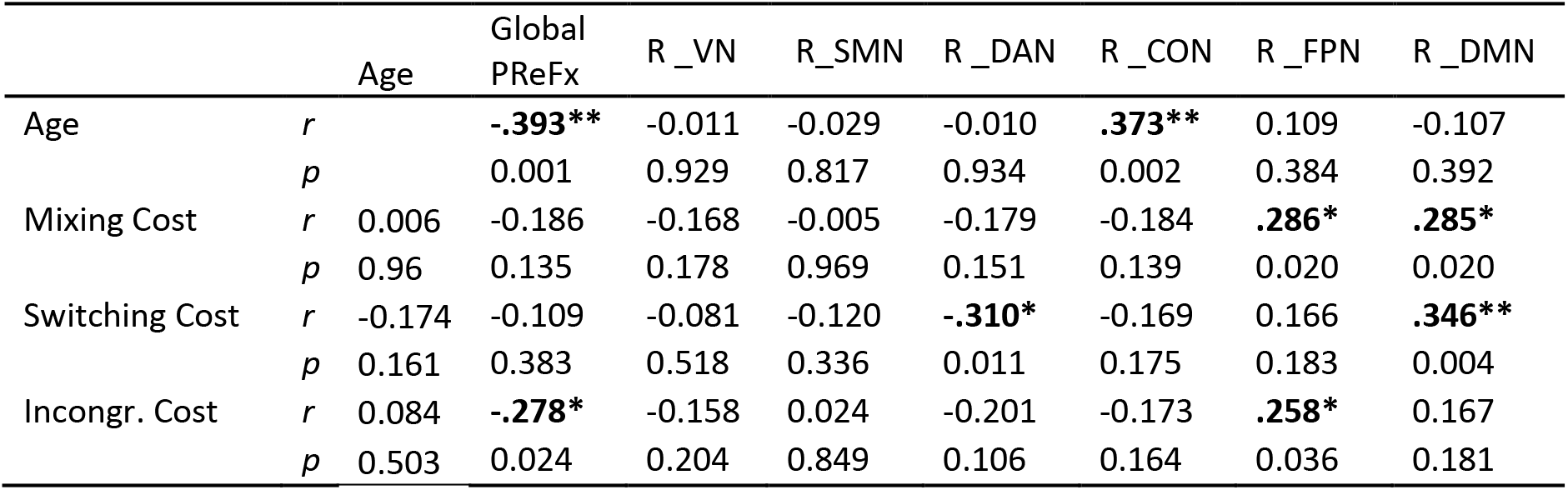
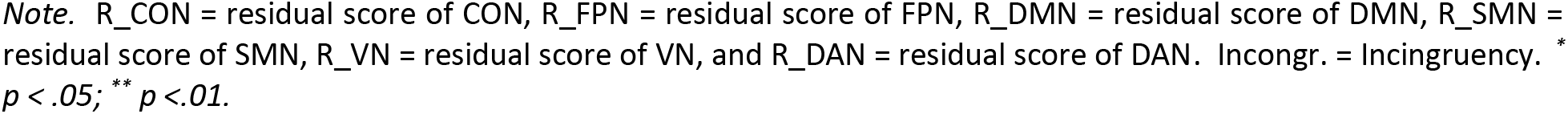
Spearman correlations between behavioral indices and global and residual PReFx values.

#### 3.2.2 Raw network PReFx

The raw network (ROI) PReFx measures were all highly correlated with each other, requiring removal of the global signal, and with incongruency cost (CON: *r* = -0.318, *p* = 0.009, DMN: *r* = -0.242, *p* = 0.044, SMN: *r* = -0.255, *p* = 0.039, VN: *r* = -0.308, *p* = 0.012, DAN: *r* = -0.311, *p* = 0.011, also see **Table S2** for the full table of results). Thus, reactive cognitive control appears to be related to the overall health of the cerebral arterial system.

#### 3.2.3 Residual network PReFx scores

As shown in **Table 3**, positive PReFx residuals in DAN were associated with smaller switching cost. In contrast, positive residuals in DMN PReFx were associated with larger mixing and switching costs. FPN’s PReFx residual values also correlated positively with mixing and incongruency costs. The positive association between age and CON’s PReFx residuals may reflect relative preservation of vascular elasticity in the CON with age, although this possibility requires further investigation.

In summary, whereas the global PReFx was most strongly associated with reactive cognitive control (incongruency cost), PReFx residuals from different networks were most strongly associated with proactive cognitive control (mixing and switching costs), indicating that proactive control may be particularly sensitive to the relative distribution of vascular elasticity across brain networks.

**Fig. 4** presents correlation maps between raw PReFx and age as well as different behavioral indices, without correction for multiple comparisons. PReFx shows a broadly distributed bilateral negative correlation with age and incongruency cost, with mixing and switching costs showing less extensive negative correlations. These negative correlations are consistent with the hypothesis that greater global arterial elasticity is associated with lower reactive control costs. For mixing and switching costs, this association is present for some brain regions and not others.

**Figure 4.**
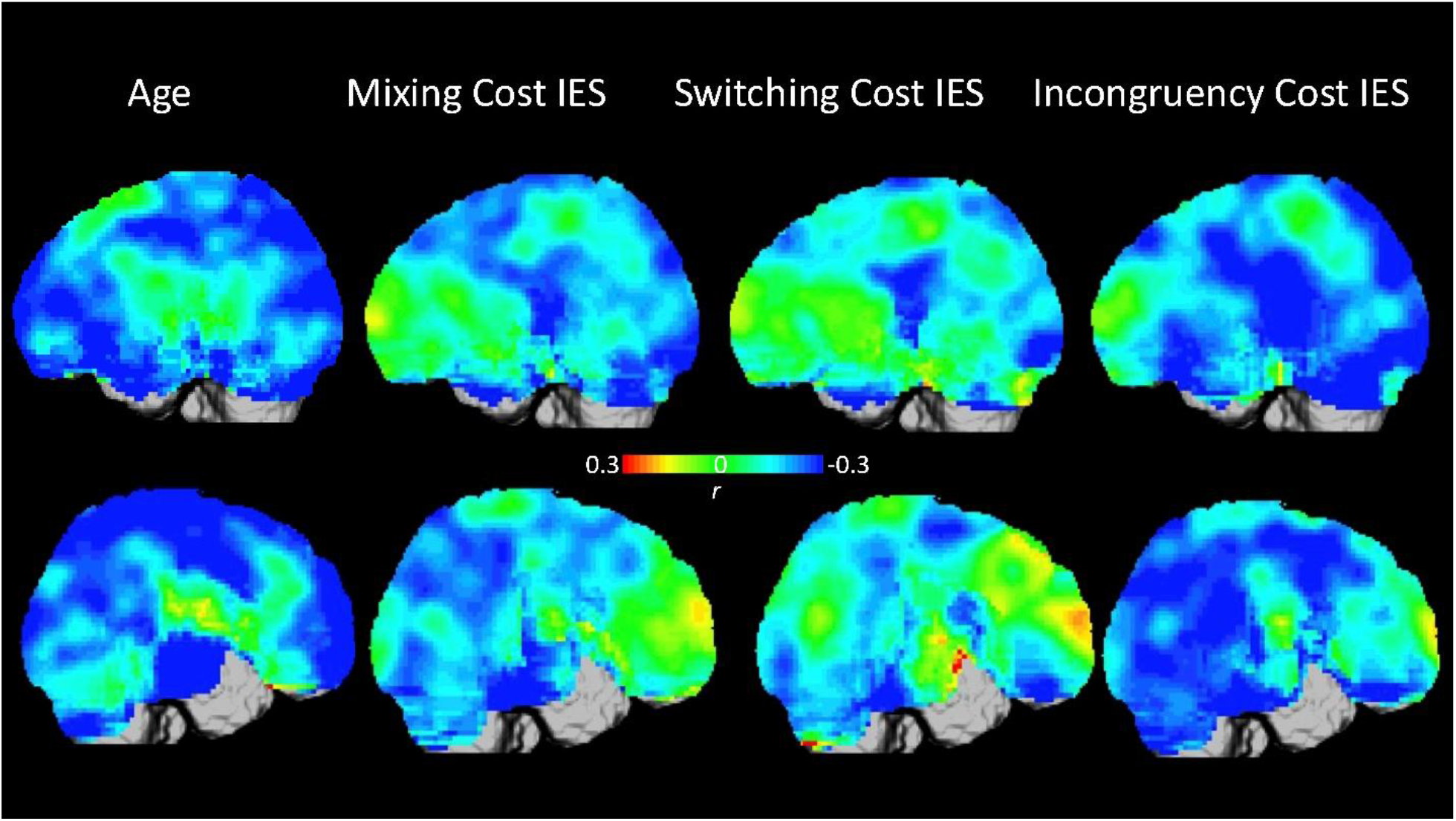
Correlation maps between raw PReFx and Age, Mixing Cost, Switching Cost, and Incongruency Cost. **Upper row**: Left sagittal view. **Bottom row**: Right sagittal view. Red colors indicate positive correlations between raw PReFx and corresponding behavioral indices, and blue colors indicate negative correlations.

### 3.3 PCA results

To better understand how the regional PReFx effects were organized, given the functional overlap of several of the networks being investigated, a PCA was conducted on the residual PReFx scores from the six brain networks (DMN, FPN, CON, DAN, SMN, and VN). **Fig. 5** summarizes the PCA results. Before rotation, the eigenvalues for the first three components were 2.60, 1.83, and 0.74 accounting for 43.27%, 30.49%, and 12.36% of the variance, respectively. After rotation, the corresponding sums of squared loadings were 2.29, 1.82, and 1.05, accounting for 38.20%, 30.35%, and 17.57% of the variance, respectively, with a cumulative variance of 86.12%. The number of components was determined based on visual inspection of the scree plot using the elbow rule (Jolliffe, 2002) and consideration of the cumulative variance explained. We decided to retain the first three components, despite the fact that the third component’s eigenvalue was < 1. In fact, in this case, the third component still accounted for a meaningful 17.57% of the total variance and the elbow rule also supported retaining the third component as shown in **Fig. 5A**.

**Figure 5.**
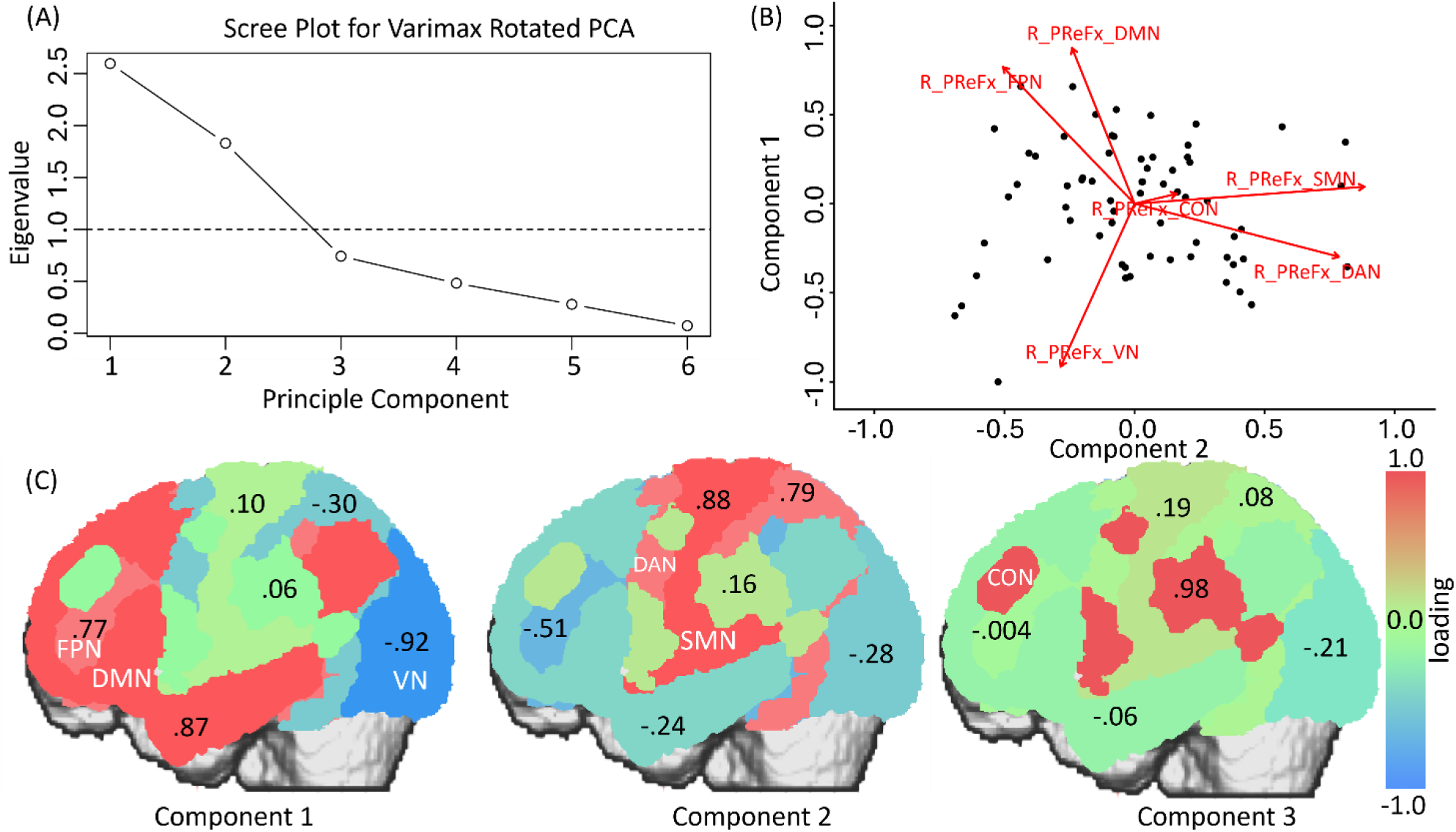
Principal component analysis of network-specific PReFx residual scores. (A) Scree plot of the component eigenvalues from the unrotated PCA. The dashed line indicates an eigenvalue of 1. (B) Biplot of components 1 and 2, showing participant component scores as points and network loading vectors as arrows. R_PReFx_CON, R_PReFx_FPN, R_PReFx_DMN, R_PReFx_SMN, R_PReFx_VN, and R_PReFx_DAN denote the PReFx deviation scores for the cingulo-opercular, frontoparietal, default mode, sensorimotor, visual, and dorsal attention networks, respectively. (C) Cortical visualization of the varimax-rotated network loadings for each component. The values displayed within each network indicate its loading on the corresponding component.

The PCA biplot (**Fig. 5B**) illustrates the distinct loading patterns of the network-specific PReFx residuals across the first two components. DMN and FPN appear to load highly on component 1, with VN loading in the opposite direction on component 1, whereas SMN and DAN loaded primarily on component 2. CON showed relatively small loadings on both components.

As shown in **Fig, 5C**, the first principal component showed high loadings for the DMN (0.874) and FPN (0.768), together with an inverse relationship to the VN (-0.915). This component therefore primarily captured shared variation in the DMN and FPN relative to the visual network. The second component was characterized by substantial loadings for DAN (0.786) and SMN (0.884). The FPN showed a loading in the opposite direction (−0.506), whereas the remaining network loadings were small. The third component exhibited a high positive loading for CON (0.983). The loadings of the other networks were small, suggesting that this component primarily reflected variation specific to the CON. **Table S3** summarized the network loadings on the three components. This analysis underscores both distinct and overlapping variance structures among these brain networks, revealing both functional commonalities and spatial proximity.

### 3.4 PLSC Results

As an exploratory analysis, PLSC was conducted to examine the multivariate relationship between PReFx residuals across the six functional networks, demographic variables (age and sex), and cognitive control performance indexed by the IES cost scores. This approach allowed for the relationships across these two blocks of variables to be evaluated jointly, thereby reducing reliance on multiple separate univariate tests. Specifically, the predictor variable block (*X*) comprised the six PReFx residual scores, along with age and sex (male=0, female=1). The dependent variable block(*Y*) included the IES for mixing cost, switching cost, and incongruency cost.

The PLSC analysis identified one significant latent pattern linking the residual PReFx scores across the functional networks to cognitive-control performance. This pattern was characterized primarily by relatively higher DAN and VN PReFx residuals and relatively lower DMN and FPN PReFx residuals in conjunction with lower mixing and switching costs. Age, sex, SMN, CON, and incongruency cost made relatively small contributions to this relationship.

As shown in **Figs. 6A** and **B**, the first latent dimension accounted for 93.26% of the cross-block covariance identified by the PLS decomposition, and the corresponding X-and Y-block latent scores were correlated at *r* = .361. A permutation test with 1000 iterations indicated that the first latent dimension was statistically significant (*p* = .023). None of the subsequent latent variables reached significance. Therefore, only the first latent variable was retained for interpretation.

**Figure 6.**
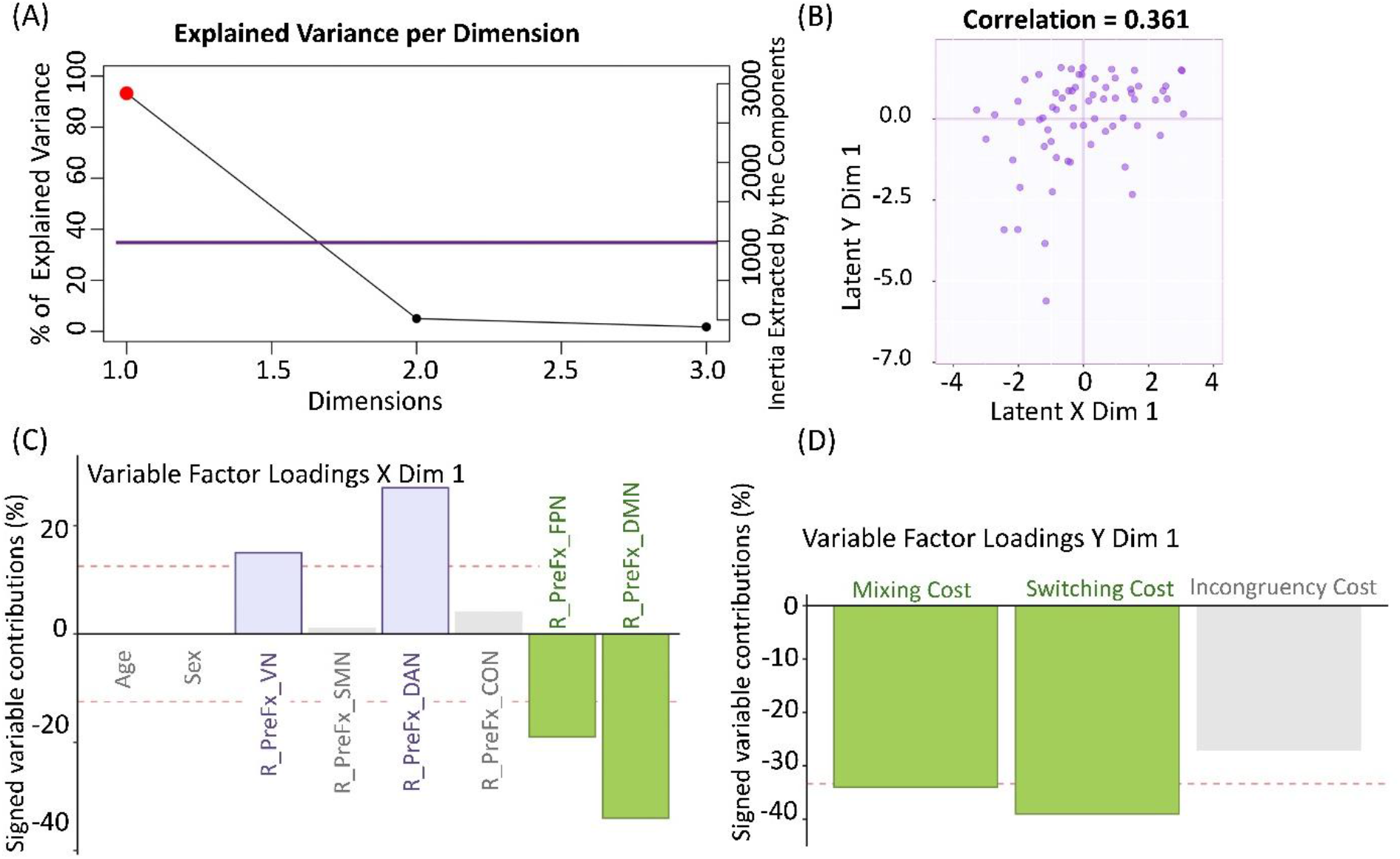
Partial least squares correlation (PLSC) results. (A) Scree plot showing the latent dimensions identified by the PLSC analysis. The purple horizontal line represents the significance threshold derived from permutation testing; dimensions exceeding this threshold were considered statistically significant. (B) Association between the X- and Y-block latent scores for the first latent dimension. (C–D) Contributions of individual variables to the X- and Y-block latent patterns, respectively, for Dimension 1. Contribution values were multiplied by 100 and rounded for visualization; their signs indicate opposite directions within the latent pattern. Red dashed lines mark the descriptive average contribution threshold for each block, calculated as 100/*p*, where *p* is the number of variables in that block.

**Fig. 6C** shows the X-block variable signed contributions for the first latent dimension. The VN and DAN contributed in one direction (purple bars, 0.152 and 0.273, respectively), whereas the DMN and FPN contributed in the opposite direction (green bars, −0.335 and −0.194). The contributions of the SMN (0.007), CON (0.0352), age (-0.0001), and sex (-0.003) were relatively small (grey bars). The stability of these contributions was evaluated using bootstrap resampling (BSR, 1000 iterations). BSR were calculated by dividing each salience by its bootstrap-estimated standard error, with absolute values of 2 or greater considered stable. DAN, FPN, and DMN met this criterion, with BSRs of 2.72, −2.66, and −3.01, respectively. Although the VN showed a relatively high contribution, its bootstrap ratio did not meet the predefined stability criterion (BSR=1.86).

**Fig. 6D** shows the Y-block variable signed contributions for the first latent dimension. Mixing, switching, and incongruency costs contributed in the same direction, with contribution values of −0.344, −0.387, and −0.269, respectively. Mixing and switching costs exceeded the average-contribution threshold, whereas incongruency cost had comparatively smaller contributions. The bootstrap analysis indicated that all three behavioral measures met the predefined stability criterion (BSR_Mixing Cost_ = -2.83, BSR_Switching Cost_ = -2.80, BSR_Incongruency Cost_ = -2.23), suggesting that their contributions were stable across bootstrap resamples.

The PLSC results reflected a multivariate pattern of covariation between network-specific PReFx residuals and cognitive-control performance. This pattern was characterized by opposing saliences for the VN/DAN and the DMN/FPN, such that relatively higher PReFx residuals in the VN and DAN, together with relatively lower residuals in the DMN and FPN, covaried with lower mixing and switching costs. These findings indicate that cognitive-control performance was related to the relative distribution of PReFx with proactive control costs. It was also compatible with the PCA results, particularly the component contrasting DMN/FPN with VN. Together, these findings provide converging evidence that mixing and switching costs are associated with a network-specific pattern of arterial elasticity, characterized by opposing contributions from DAN/VN and DMN/FPN.

## 4. DISCUSSION

This study employed a network-based approach to examine the relationship between cerebrovascular status (indexed by PReFx pulse-DOT measures) and cognitive control performance in a task-switching paradigm that allows for separation of proactive and reactive control processes. Consistent with previous studies using this paradigm, participants exhibited reliable mixing, switching, and incongruency costs (Karayanidis et al., 2011). There was also a main effect of age on cost effects, but no evidence that the magnitude of the mixing, switching, or incongruency costs interacted with age. At the *global PReFx* level, we first replicated the presence of age-related decline in global vascular elasticity (Zimmerman et al., 2021). We then found that higher global PReFx (i.e. better arterial elasticity) was associated with better reactive control (as indexed by lower incongruency cost), but not with proactive control. These same patterns were also observed with *raw network PReFx* analyses, potentially reflecting the strong shared influence of the global signal. *Residual network PReFx* values were therefore calculated by removing the global signal to enhance sensitivity to network-specific variations. PCA results revealed the spatial organization of the residual network PReFx: three components primarily characterized by (1) the DMN and FPN (with opposing VN loadings), (2) the DAN and SMN, and (3) the CON. Finally, the PLSC analysis revealed a significant multivariate relationship between the residual network PReFx scores and proactive control costs, capturing an opposing pattern of network-specific PReFx deviations: Higher PReFx residuals in VN and DAN, together with lower PReFx residuals in DMN and FPN, covaried with lower proactive control costs (mixing and switching cost). Together, these converging results suggest that reactive control is more closely associated with global cerebrovascular elasticity, whereas proactive control is related to the *relative* distribution of cerebrovascular elasticity across functional networks.

### 4.1 Cerebrovascular network organization: PCA analyses

The first principal component was primarily represented by the DMN and FPN, with the VN showing an opposing pattern. This pattern may reflect shared cerebrovascular variation across the DMN and FPN, which is distinct from the VN. The DMN is often characterized as a task-negative network, known for its role in self-referential and internally directed thought processes (Xia et al., 2022). The FPN, while traditionally characterized as a task-positive network involved in cognitive control and higher-order processing, shares a close spatial proximity with the DMN, and exerts a strong role in top-down inhibitory control. Their clustering in the PReFx residual analysis may partly reflect their spatial proximity, as illustrated in **Fig. 5C**, which could contribute to their shared variance. This may also reflect the relatively limited spatial resolution of cerebrovascular phenomena relative to those underlying functional connectivity. In contrast, VN loaded strongly in the opposite direction to the DMN and FPN, indicating that this component primarily captured a contrast between PReFx variation in the DMN/FPN and the VN.

The second principal component was primarily represented by the DAN and SMN, suggesting a shared pattern of cerebrovascular variation across these networks. DAN is primarily responsible for top-down attention control, directing focus to task-relevant stimuli (Corbetta & Shulman, 2002). SMN, on the other hand, is involved in motor execution and sensory processing, translating sensory input into physical responses (Yeo et al., 2011). As for FPN and DMN, their clustering may result from their spatial proximity (see **Fig. 5C**), and/or functional similarities, as both networks are activated during task-related processes, contributing to cognitive control through attention and motor coordination (Cocchi et al., 2013).

The third principal component was primarily represented by the CON. Given the CON’s role in cognitive control and salience detection, its separation from the other networks may suggest a relatively distinct pattern of cerebrovascular variation within this network. However, it is important to note that the PCA structure should be interpreted in light of the limited spatial sensitivity of the optical coverage as discussed in the introduction. As Pulse-DOT primarily samples cortical regions, networks with deeper structures, such as the CON, may be less completely captured.

### 4.2 Dual Mechanisms of Cognitive Control and Cerebrovascular Status

The *global* PReFx was negatively correlated with reactive control cost, even after controlling for age and sex, which suggested that when the brain as a whole is exhibits greater cerebrovascular elasticity, all network regions are likely to have the necessary blood flow and therefore better equipped to react quickly to resolve immediate cognitive conflict. As a result, individuals with higher global PReFx are better at managing reactive control. In contrast, *residual* PReFx scores, which reflect how much specific networks deviate from the global cerebrovascular elasticity signal, were more closely associated with proactive control measures, with positive residuals in DAN’s PReFx showing a negative relationship with mixing and switching costs and positive residuals in DMN and FPN’s PReFx showing a positive relationship with these costs. These associations are broadly consistent with the task-positive role of the DAN and the task-negative role of the DMN. The FPN pattern was less expected and will be further discussed in the next section (4.3).

Thus, these results suggest that variations in the elasticity of the cerebrovascular system, both globally and regionally, may exert distinct influences on the two types of cognitive control: better global cerebrovascular status was associated with smaller incongruency costs, suggesting a relationship with target-triggered interference resolution. In contrast, proactive control was associated with the relative configuration of PReFx across functional networks.

### 4.3 Balance between the cerebrovascular system and cognitive control performance

The PLSC analysis provides a nuanced understanding of how network-specific PReFx deviations from the global signal covary with cognitive-control performance. Specifically, DMN and FPN displayed a positive relationship with mixing and switching costs, while VN and DAN showed an opposite relationship. As the residual PReFx scores quantify network-specific deviations from global arterial elasticity, they provide an index of the *relative* balance of cerebrovascular elasticity across functional networks. The PLSC results further indicate that relative balance in the cerebrovascular system may be relevant for flexible and efficient cognitive control processes.

Greater PReFx residuals in the DMN and FPN were found to correlate with poorer proactive control, as indicated by larger costs in the PLSC results. This suggests that cerebrovascular irregularities within these networks may compromise the brain’s capacity to engage effectively in anticipatory, goal-directed processing. The DMN, a task-negative network, typically deactivates during goal-oriented tasks and may be particularly susceptible to this disruption. When vascular elasticity within the DMN exceeds the global average, it is plausible that this network becomes more predisposed to facilitating non-task-related activities, such as mind-wandering or introspection (Anticevic et al., 2012; Raichle, 2015), rather than supporting sustained cognitive engagement, resulting in increased proactive control costs. The role of the FPN is less clear, but it may be largely driven by its spatial proximity to the DMN rather than a direct functional impairment of cognitive control, as the DMN showed a larger effect compared to the FPN in both the correlation and PLSC analyses. This spatial proximity assumption is also supported by the PCA analysis, where DMN and FPN formed a component. As illustrated in **Fig. 4C**, the vascular territories encompassing the FPN and DMN are spatially adjacent.

Conversely, greater residuals in the DAN and VN were associated with better proactive control, which suggests that these networks may function more effectively in supporting sustained, goal-directed processes when their vascular states are more enhanced compared to other regions. The DAN is involved in top-down, goal-directed attention, which is essential for maintaining proactive control (Vossel et al., 2014), and the VN is essential for visual processing. These networks may benefit from greater vascular adaptability, allowing them to efficiently engage in visual proactive control tasks. This contrast between task-positive and task-negative networks indicates that proactive control may be strongly influenced by how well the brain balances resource allocation between these competing networks.

A similar conceptual parallel of balancing mechanism could be found in prior work suggesting that adaptive control depends on coordinated contributions from functionally distinct, and sometimes opposing, brain systems. For example, studies of cognitive–affective interactions have supported a dissociated role of dorsal “cold” executive and ventral “hot” affective systems in supporting goal-directed behavior (Dolcos et al., 2008; Iordan et al., 2013). Although the dorsal “cold” and ventral “hot” systems do not map directly onto the functional networks examined here, they show partial functional and anatomical overlap with these networks, providing a conceptual parallel for understanding cognitive control in a perspective of coordinated balance among distributed brain systems.

The degree of variation in cerebrovascular elasticity across these networks may also represent a neural marker of how well the brain manages this balancing act. When networks like DAN are well-supplied and functional, the brain can better maintain proactive control. Conversely, when networks associated with internally directed and affectively relevant processing, such as the DMN, are relatively better supported vascularly, this may lead to resource misallocation in cognitive control, impairing the brain’s ability to maintain task goals over time. These results highlight a nuanced balance between global and regional cerebrovascular function and suggest that cognitive control may hinge on the relative “fitness” of these networks’ vascular health. Maintaining an equilibrium, where global PReFx is optimally configured, but regional networks can deviate in ways that support specific cognitive demands, may represent an adaptive cerebrovascular state. Disruption to this balance, as seen in excessive variations within the DMN, could lead to increased cognitive costs in proactive control.

### 4.4 Cerebrovascular age and chronological age

In this study, chronological age was significantly associated with lower global PReFx, replicating previous findings that cerebral arterial elasticity declines across adulthood (Fabiani et al., 2014; Tan et al., 2019; Bowie et al., 2024). This relationship was broadly distributed across the cortical mantle, suggesting that age-related cerebrovascular changes were not restricted to a single functional network. In contrast, chronological age was not significantly associated with mixing, switching, or incongruency costs. Thus, although the sample showed clear age-related variations in cerebrovascular status, these differences were not accompanied by a simple linear decline in the cognitive control indices examined here, consistent with prior work showing heterogeneous age effects across task-switching measures (Karayanidis et al., 2011).

Several factors may help explain this dissociation. First, the sample consisted of healthy, educated adults, which may have reduced the expression of age-related differences in cognitive-control performance through greater cognitive reserve (Stern et al., 2023). Second, the relationship between aging and cognitive control is heterogeneous: individuals of the same chronological age can differ considerably in vascular health, general cognitive abilities, and the strategies they use to perform the same task. Chronological age may therefore be too coarse a marker to capture the biological and functional heterogeneity underlying individual differences in performance. At the same time, our findings do not indicate that age was irrelevant: Age was associated with global PReFx decline and deviation in the CON. However, age and sex contributed minimally to the significant PLSC pattern linking network-specific PReFx deviations with cognitive-control costs. This suggests that the multivariate network pattern was not simply a reflection of participants’ chronological age. Instead, individual differences in the relative distribution of cerebrovascular elasticity across networks appeared to provide information about cognitive-control performance that was not captured by age alone. Thus, although chronological age was not associated with cognitive-control costs, PReFx captured global and network-specific cerebrovascular variation that was related to performance, highlighting its potential value as a more functionally informative marker than age alone.

### 4.5 Limitations and future direction

It is important to note that diffuse optical imaging is limited in its ability to measure vascular activity in deeper brain regions. Pulse-DOT predominantly captures superficial cortical layers and cannot capture the contribution of deeper structures—including subcortical regions like the hippocampus, amygdala and thalamus. However, the major cerebral arteries largely course along the cortical surface and are therefore accessible to optical measurements. In addition, previous work has shown that PReFx is sensitive to downstream age-related differences in brain structure and function, as well as cognitive performance (Bowie et al., 2024; Tan et al., 2017; Kong et al., 2020; Clements et al., 2025). Thus, although pulse-DOT cannot directly characterize deeper brain structures, PReFx may still provide information relevant to broader brain and cognitive outcomes associated with cerebrovascular health.

To overcome these constraints, future research could benefit from employing multimodal imaging approaches, such as integrating optical imaging with fMRI (Blood Oxygenation Level Dependent or Arterial Spin Labeling), or new approaches based on fMRI (Marchini et al., 2026). This combination would enable the capture of both surface-level and deeper vascular dynamics, providing a more comprehensive mapping of cerebrovascular function.

Furthermore, while this study focused on cognitive control tasks, extending the investigation to include affective and memory tasks may provide a more comprehensive understanding of the cerebrovascular balance mechanism. Previous research has shown that the dorsal “cold” executive system and the ventral “hot” affective system interact to balance cognitive and emotional functions. Future research could explore whether specific cerebrovascular configurations differentially predict performance on cognitive versus emotional tasks. This line of inquiry may help refine our understanding of the distinct roles that cerebrovascular health and elasticity play across various functional domains, enhancing our knowledge of brain network dynamics in both cognitive and affective contexts.

### 4.6 Conclusions

The present study introduces a network-based approach to the study of cerebrovascular health by examining the intrinsic organization of vascular elasticity across large-scale functional networks and its association with cognitive control performance. Previous work has largely viewed cerebrovascular health as a global property, thereby overlooking the intrinsic organization of the cerebrovascular system. By analyzing residual PReFx across functional networks, the present study showed that network-specific vascular elasticity was organized into coherent components, including a DMN-FPN component with the VN loading in the opposite direction, a DAN-SMN component, and a CON-dominated component. This organization may reflect both spatial proximity among vascular territories and functional relatedness among large-scale brain networks. Importantly, residual PReFx across different networks was differentially associated with proactive control costs. Relatively greater residual PReFx in the VN and DAN was associated with better proactive control, whereas relatively greater residual PReFx in the DMN and FPN was associated with poorer proactive control. In contrast, reactive-control performance was associated primarily with higher PReFx in global and raw PReFx in all networks. These findings suggest that proactive control may be related more closely to the relative distribution of cerebrovascular elasticity across networks, whereas reactive control may depend more on shared global cerebrovascular status. They also indicate that cerebrovascular health is not only a global index, but a spatially organized network-level property. The relative balance of vascular elasticity across functional networks may contribute to individual differences in cognitive control performance.

## Acknowledgments

This research was supported by NIA grants R01AG059878 and RF1AG062666 to M. Fabiani and G. Gratton. We acknowledge the support of the Bioimaging Center of the Beckman Institute for Advanced Science and Technology at the University of Illinois Urbana-Champaign (UIUC-BI-BIC).

