## Supplemental figures and tables for "Network-related variations of cerebrovascular status are correlated with cognitive control performance"

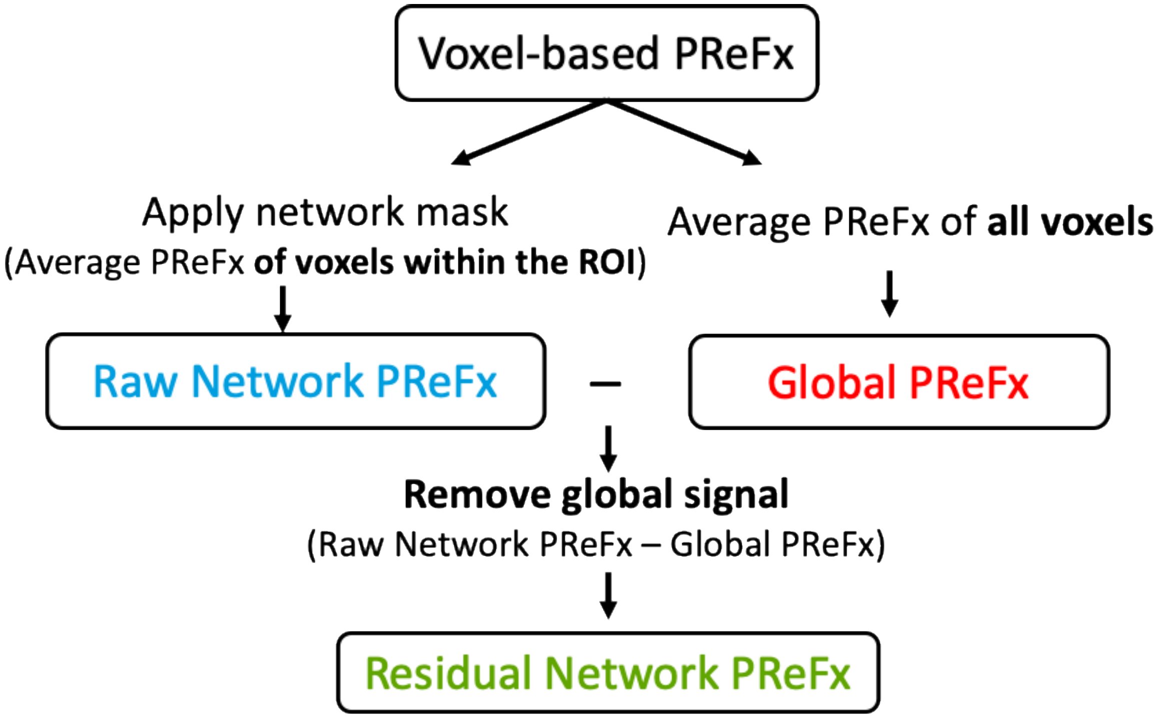


**Figure S1.** Flowchart illustrating the computation of Global, Raw Network, and Residual Network PReFx. Voxel-based PReFx maps were first generated using Opt3D software (Gratton, 2000). Global PReFx was calculated by averaging PReFx values across all voxels, as described in the main text. Raw network PReFx was obtained by applying network ROI masks and averaging voxel PReFx values within each ROI. Finally, the global PReFx signal was subtracted from each network-based PReFx to derive the residual network PReFx.

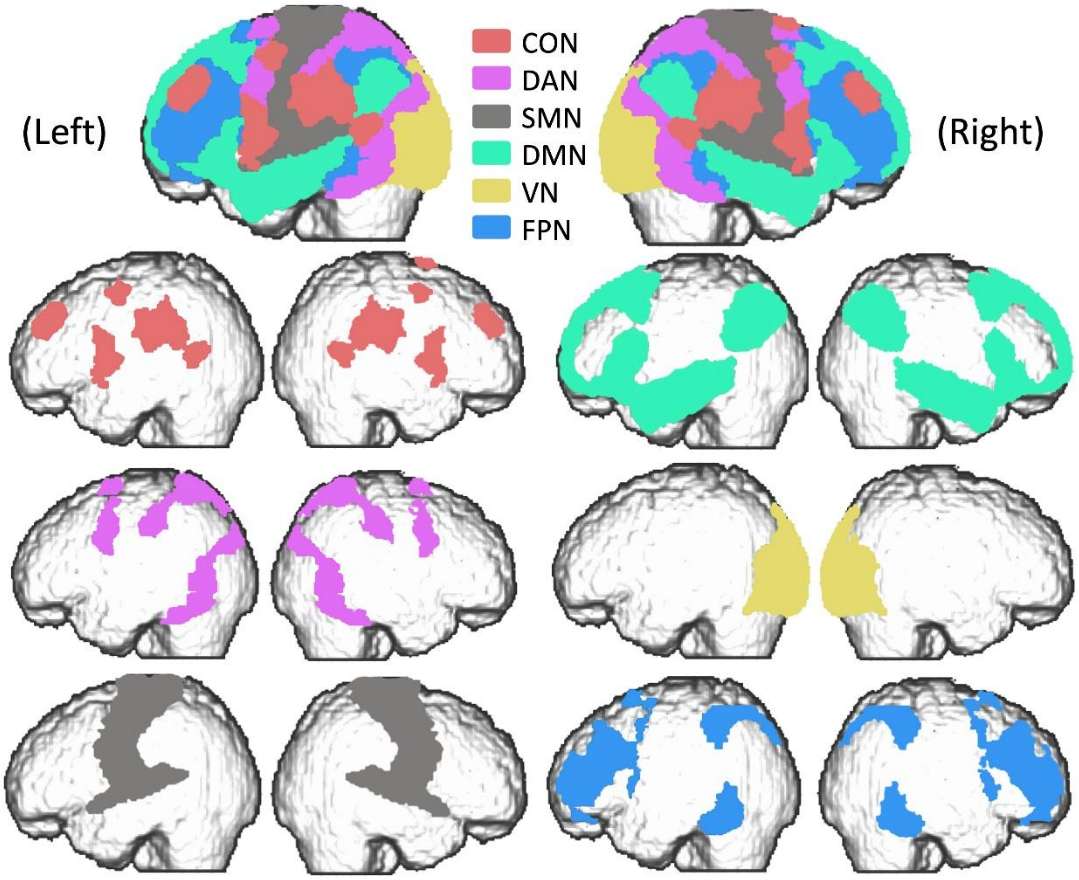


**Figure S2** Cortical surface visualization of the six functional network ROIs based on the Yeo et al. (2011) atlas. The top panels show the six network ROIs overlaid on the cortical surface in the following order, from the topmost to the bottommost layer: Cingulo-Opercular Network (CON), Dorsal Attention Network (DAN), Somatomotor Network (SMN), Visual Network (VN), Default Mode Network (DMN), and Frontoparietal Network (FPN). The lower panels show each network separately. In each pair of images, the left and right panels represent sagittal views of the left and right hemispheres, respectively.

**Table S1.** Full results of linear mixed-effects models for reaction time (RT) and accuracy (ACC) across mixing, switching, and incongruency costs with age as a predictor.

| Outcome | Effect | Estimate | SE | *df* | *t* | *p* |
|---|---|---|---|---|---|---|
| Mixing cost | **Dependent variable: RT** | | | | | |
| | Age | 2.71 | 1.01 | 64 | 2.69 | 0.009* |
| | Trial type (AR, MR) | 79.04 | 10.86 | 64 | 7.28 | < 0.001*** |
| | Age × Trial type | 0.22 | 0.75 | 64 | 0.30 | 0.765 |
| | **Dependent variable: ACC** | | | | | |
| | Age | 0.02 | 0.01 | 64 | 1.86 | 0.068 |
| | Trial type (AR, MR) | -0.22 | 0.11 | 64 | -2.01 | 0.049* |
| | Age × Trial type | 0.01 | 0.01 | 64 | 1.42 | 0.160 |
| Switching cost | **Dependent variable: RT** | | | | | |
| | Age | 2.52 | 1.53 | 64 | 1.65 | 0.103 |
| | Trial type (MR, MS) | 64.76 | 9.56 | 64 | 6.77 | < 0.001*** |
| | Age × Trial type | -0.61 | 0.66 | 64 | -0.93 | 0.358 |
| | **Dependent variable: ACC** | | | | | |
| | Age | 0.20 | 0.01 | 64 | 1.83 | 0.072 |
| | Trial type (AR, MR) | -0.59 | 0.10 | 64 | -6.24 | < 0.001*** |
| | Age × Trial type | -0.01 | 0.01 | 64<br>4 | -1.59 | 0.1174 |
| Incongruency cost | **Dependent variable: RT** | | | | | |
| | Age | 2.59 | 1.17 | 64 | 2.21 | 0.031* |
| | Trial type (AR, MR) | 85.81 | 8.48 | 64 | 10.12 | < 0.001*** |
| | Age × Trial type | 0.47 | 0.58 | 64 | 0.81 | 0.424 |
| | **Dependent variable: ACC** | | | | | |
| | Age | 0.01 | 0.01 | 64 | 1.53 | 0.132 |
| | Trial type (AR, MR) | -0.79 | 0.09 | 64 | -8.56 | < 0.001*** |
| | Age × Trial type | -0.01 | 0.01 | 64 | -1.10 | 0.277 |

*Note.* Degrees of freedom were approximated using Satterthwaite's method. ACC was logit-transformed when modeled as the dependent variable to reduce skewness associated with ceiling-level performance. Values of 0 and 1 were bounded to 0.001 and 0.999 before logit transformation. * $p < .05$; ** $p < .01$; *** $p < 0.001$

**Table S2**. Full uncorrected Spearman correlation table between cognitive costs and raw network PReFx.

| | | CON | FPN | DMN | SMN | VN | DAN | Global |
|---|---|---|---|---|---|---|---|---|
| Age | *r* | -.264* | -.353** | -.411** | -.348** | -.315** | -.368** | -.393** |
| | *p* | 0.032 | 0.004 | <.001 | 0.004 | 0.010 | 0.002 | 0.001 |
| MC_RT | *r* | -.288* | -0.159 | -0.157 | -0.231 | -.276* | -.273* | -0.222 |
| | *p* | 0.019 | 0.202 | 0.209 | 0.062 | 0.025 | 0.027 | 0.073 |
| SC_RT | *r* | -0.156 | -0.024 | 0.005 | -0.169 | -0.091 | -0.169 | -0.081 |
| | *p* | 0.210 | 0.847 | 0.968 | 0.174 | 0.465 | 0.174 | 0.519 |
| MC_ACC | *r* | -0.017 | 0.046 | 0.046 | 0.045 | -0.020 | -0.036 | 0.023 |
| | *p* | 0.893 | 0.716 | 0.715 | 0.719 | 0.871 | 0.772 | 0.855 |
| SC_ACC | *r* | -0.080 | -0.072 | -0.015 | -0.067 | -0.020 | -0.053 | -0.066 |
| | *p* | 0.525 | 0.568 | 0.908 | 0.595 | 0.875 | 0.672 | 0.601 |
| MC_IES | *r* | -.256* | -0.123 | -0.115 | -0.183 | -.249* | -.247* | -0.186 |
| | *p* | 0.038 | 0.327 | 0.356 | 0.141 | 0.044 | 0.046 | 0.135 |
| SC_IES | *r* | -0.173 | -0.06 | -0.019 | -0.167 | -0.118 | -0.179 | -0.109 |
| | *p* | 0.165 | 0.635 | 0.877 | 0.179 | 0.346 | 0.151 | 0.383 |
| IC_RT | *r* | -.336** | -0.229 | -.272* | -.277* | -.294* | -.323** | -.283* |
| | *p* | 0.006 | 0.064 | 0.027 | 0.024 | 0.016 | 0.008 | 0.021 |
| IC_ACC | *r* | -0.067 | 0.027 | 0.013 | -0.017 | -0.074 | -0.069 | -0.043 |
| | *p* | 0.591 | 0.831 | 0.919 | 0.892 | 0.557 | 0.582 | 0.733 |
| IC_IES | *r* | -.318** | -0.211 | -.242* | -.255* | -.308* | -.311* | -.278* |
| | *p* | 0.009 | 0.090 | 0.044 | 0.039 | 0.012 | 0.011 | 0.024 |
| CON | *r* | 1.00 | .872** | .871** | .888** | .825** | .921** | .922** |
| | *p* | . | <.001 | <.001 | <.001 | <.001 | <.001 | <.001 |
| FPN | *r* | .872** | 1.00 | .976** | .836** | .831** | .890** | .961** |
| | *p* | <.001 | . | <.001 | <.001 | <.001 | <.001 | <.001 |
| DMN | *r* | .871** | .976** | 1.00 | .849** | .834** | .908** | .967** |
| | *p* | <.001 | <.001 | . | <.001 | <.001 | <.001 | <.001 |
| SMN | *r* | .888** | .836** | .849** | 1.00 | .785** | .927** | .902** |
| | *p* | <.001 | <.001 | <.001 | . | <.001 | <.001 | <.001 |
| VN | *r* | .825** | .831** | .834** | .785** | 1.00 | .898** | .922** |
| | *p* | <.001 | <.001 | <.001 | <.001 | . | <.001 | <.001 |
| DAN | *r* | .921** | .890** | .908** | .927** | .898** | 1.00 | .963** |
| | *p* | <.001 | <.001 | <.001 | <.001 | <.001 | . | <.001 |
| Global | *r* | .922** | .961** | .967** | .902** | .922** | .963** | 1.00 |
| | *p* | <.001 | <.001 | <.001 | <.001 | <.001 | <.001 | . |

Note. MC = Mixing Cost; SW = Switching Cost; IC = Incongruency Cost; CON = Cingulo-Opercular Network; DAN = Dorsal Attention Network; SMN = Somatomotor Network; VN = Visual Network; DMN = Default Mode Network; FPN = Frontoparietal Network. * $p < .05$; ** $p < .01$.

**Table S3.** Varimax-Rotated Network Loadings on the Principal Components.

| Network | RC1 | RC2 | RC3 |
|---|---|---|---|
| R_PReFx_VN | -0.915 | -0.284 | -0.208 |
| R_PReFx_SMN | 0.095 | 0.884 | 0.191 |
| R_PReFx_DAN | -0.298 | 0.786 | 0.075 |
| R_PReFx_CON | 0.056 | 0.162 | 0.983 |
| R_PReFx_FPN | 0.768 | -0.506 | -0.004 |
| R_PReFx_DMN | 0.874 | -0.24 | -0.059 |

*Note*. RC = rotated component; R_PReFx = residual network PReFx; VN = Visual Network; SMN = Somatomotor Network; DAN = Dorsal Attention Network; CON = Cingulo-Opercular Network; FPN = Frontoparietal Network; DMN = Default Mode Network. Values are varimax-rotated component loadings from the principal component analysis.
